# Leveraging AlphaFold 3 for Structural Modeling of Neurological Disorder-Associated Proteins: A Pathway to Precision Medicine

**DOI:** 10.1101/2024.11.18.624211

**Authors:** Nishant Gadde, Sachi Dodamani, Rayaan Altaf, Sanjit Kumar

## Abstract

Accurate structural modeling of neurological disorder-causing proteins provides an important layer in unraveling the mechanism of disease and identifying therapeutic targets. This study utilizes AlphaFold 3, a state-of-the-art protein structure prediction platform, to model and interpret cis- and trans-pQTL-derived proteins associated with Alzheimer’s disease, Parkinson’s disease, and stroke. Using the NG00102 dataset, we created a high-resolution structure for more than 1,200 proteins expressed in Brain, CSF, and Plasma, providing tissue-specific protein structure analysis with associated functional implications. AlphaFold 3 predictions have illuminated key structure parameters including sequence length, average pLDDT confidence scores, and overall distribution of residues with confidence of >75% pLDDT. We used these features to determine the set of druggable proteins having optimal sequence lengths of 100-3000 residues, high structural reliability as evidenced by an average pLDDT > 80, and contain large regions of high-confidence residues. Tissue-specific mapping revealed unique mechanisms characterized by both cis and trans-pQTL effects, that have critical functional implications for how these genetic variants act in neurological disease pathways. Protein clusters by structural properties then led to more defined subgroups with potential implications for drug intervention. This integrated effort captures the strength of AlphaFold 3 in linking genetic variation to protein structure and function, providing a scalable pipeline for prioritizing therapeutic targets. Coupling our results with advanced predictive modeling and tissue-specific data sets provides a robust framework for uncovering new mechanisms and druggable targets in the research of Alzheimer’s, Parkinson’s, and stroke. This advances the field toward precision medicine.

## Introduction

Neurological disorders, including Alzheimer’s disease, Parkinson’s disease, and stroke, are complex and multifactors that have strong genetic contributions. Of these, pQTLs represent a potent strategy for genetically linking variants into protein expression levels and insight into tissues of specific mechanisms in disease. However, the detailed understanding of the structural and functional characterization of the effect of both cis- and trans-pQTLs on protein functions has become a big bottleneck in translation from genetic data to therapeutic targets. It has been shown that genetic variants are crucially influencing the expression pattern of proteins in CSF, hence indicating neurodegenerative diseases such as Alzheimer’s and Parkinson’s diseases (Sasayama et al., 2016).

High-resolution modeling of protein structure has revolutionized with the advancement of computational biology. AlphaFold 3, the latest iteration of DeepMind’s AlphaFold technology, represents a transformative tool in structural biology. It enables accurate, large-scale prediction of protein structures with confidence metrics, such as the per-residue pLDDT score, which can be used to prioritize high-confidence regions for downstream analysis. AlphaFold has demonstrated substantial potential in elucidating protein interactions and identifying novel drug targets in diseases like Alzheimer’s and stroke (Liu et al., 2023).

The NG00102 dataset offers a unique opportunity to investigate structural and functional consequences of the pQTLs in neurodegenerative diseases of Alzheimer’s, Parkinson’s, and stroke. The dataset involves proteins expressed in tissues highly relevant to neurological diseases, including the brain, cerebrospinal fluid, and plasma, along with comprehensive genetic data. Recent work mapped thousands of proteins in plasma and CSF for genetic associations with diseases, including Alzheimer’s and cardiovascular conditions (Sun et al., 2022). Proteomic investigations that integrate data across multiple tissues have recently pointed out that combining pQTL mapping with structural predictions is essential for tissue-specific mechanisms (Yang et al., 2020).

In this study, we extend AlphaFold 3 to model and interpret the structures of more than 1,200 pQTL-derived proteins from the NG00102 dataset. Using a pipeline that integrates sequence analysis, structural modeling, and confidence metrics, we identify druggable proteins based on their structural and functional parameters; chart tissue-specific mechanisms influenced by both cis- and trans-pQTLs according to Fauman & Hyde (2022); and prioritize proteins for therapeutic exploration against AD, PD, and stroke. Our results now not only help bridge the gap between genetic variation and protein function but also help build a burgeoning framework for precision medicine in neurological research. This is one example of how AlphaFold 3 could help integrate genetic and structural data to understand the pathogenesis of a complex disease.

## Literature Review

### Introduction to Protein Structure Prediction

Protein structure prediction is one of the most aggressively pursued areas in bioinformatics. Applications range from drug discovery to unraveling disease mechanisms to the elucidation of biological pathways. Traditional methods of X-ray crystallography, nuclear magnetic resonance spectroscopy, and cryo-electron microscopy have been highly instructive in providing insights into protein structures. However, these methodologies are resource-intensive and generally time-consuming for large-scale studies. Recent breakthroughs introduced deep learning-based tools such as AlphaFold, which have revolutionized the pace at which structural biology is done by predicting protein structures at a near-experimental level of accuracy (Jumper et al., 2021).

### AlphaFold 3 and Capabilities

AlphaFold 3 now extends the capability of its forerunners to include multi-chain interaction predictions while further expanding confidence metrics such as the per-residue predicted LDDT score. These advancements enable detailed annotations of biomolecular interactions, which are crucial for understanding cis- and trans-pQTL effects in disease-relevant proteins (Liu et al., 2023). By leveraging AlphaFold 3, researchers can now assess not only the static structure of individual proteins but also their potential interactions and conformational changes, which are essential for understanding complex diseases like Alzheimer’s and Parkinson’s.

### pQTL Studies in Neurological Disorders

Protein quantitative trait loci are genetic variants that influence protein abundance. They constitute a critical link between genotype and phenotype and, for this reason, are useful in deciphering the molecular basis of diseases. While cis-pQTLs result in altered expression of proteins as a consequence of variants near the encoding gene, trans-pQTLs influence proteins encoded by distant loci through regulatory networks. The studies have discovered the importance of pQTL mapping in CSF, plasma, and brain tissues in providing insight into the pathophysiology of AD, PD, and stroke (Sasayama et al., 2016; Sun et al., 2022).

### Integration of pQTL Data with Structural Models

The combination of multi-tissue proteomic data with high-resolution genetic mapping in the NG00102 dataset offers a resource for studying pQTL effects in neurological disorders without parallel. AlphaFold 3 further enhances this dataset by providing high-confidence structural predictions for proteins influenced by cis- and trans-pQTLs. Integrative studies combining pQTLs data with AlphaFold models have illustrated the potential to identify disease-specific protein conformations and druggable regions, as illustrated by Yang et al., 2020.

### Therapeutic Target Prioritization

One of the most significant challenges in drug discovery is identifying druggable proteins. The structural predictions from AlphaFold 3 will inform the prioritization of proteins with high-confidence, stable regions, together with pQTL data. In particular, proteins with a sequence length in the range of 100-3000 residues, pLDDT scores >80, and high-confidence residues (>75%) are well-suited for drug targeting. This approach has been applied to prioritize therapeutic targets in Alzheimer’s and Parkinson’s research, with significant findings linking structural features to tissue-specific expression and disease mechanisms (Fauman & Hyde, 2022).

## Methodology

Using AlphaFold 3 predictions and the NG00102 dataset, we propose an integrative workflow that allows for the investigation of the structural and functional implications of both the cis- and trans-pQTL-derived proteins in AD, PD, and stroke. Below, in paragraphs, is a detailed methodology, along with specifications regarding the placement of visual elements.

### 1. Data Acquisition

The NG00102 dataset, obtained from the Data Sharing Service (DSS) of Niagads, represents a high-resolution investigation of protein quantitative trait loci (QTLs). It focuses on proteins expressed in key tissues—brain, cerebrospinal fluid (CSF), and plasma—each critically implicated in the pathophysiology of Alzheimer’s disease, Parkinson’s disease, and stroke. This dataset fuses genetic variability with proteomic data, leading to singular insight into the ways genetic variability affects protein expression and function in a tissue-specific manner. This dual-layered integration of genetic and proteomic information hereby positions the NG00102 dataset as an extremely valuable resource for structural and functional inquiries, particularly for pinpointing the pathways and targets of interest that are tissue-specific and druggable in neurodegenerative diseases.

### 2. Data Preparation

The identifiers of proteins, generally UniProt IDs, were retrieved from the NG00102 dataset. These identifiers were segregated based on tissue of origin: brain, CSF, and plasma. This was done to ensure that the analysis was tissue-specific. The list was further filtered to identify proteins with significant tissue-specific expression that might be associated with neurological disorders. For example, proteins with high expression in CSF are more likely to affect cerebrovascular conditions, while brain-specific proteins may directly influence Alzheimer’s and Parkinson’s disease pathways. These tissue-specific lists of proteins were curated into input for structural modeling, ensuring that only biologically relevant candidates were targeted in the downstream analyses.

### 3. AlphaFold 3 Structural Prediction

AlphaFold 3 is the latest in DeepMind’s suite of deep-learning-based tools for protein structure prediction. Deployed in a Google Colab environment, scripts were executed to model the structures of over 1,200 proteins, delivering residue-level confidence metrics as pLDDT scores. The protein sequences were formatted into FASTA files and submitted to the AlphaFold 3 server for high-resolution structural modeling. Outputs of predictions were in PDB format, including 3D structures with per-residue confidence metrics. This added much robustness to the prediction pipeline and ensured that only quality structural models would be produced for further analysis, having the pLDDT metrics act as a critical benchmark in evaluating the reliability of each prediction.

### 4. Structural Model Analysis

AlphaFold-generated PDBs were analyzed using multiple parameters for filtering proteins regarding their potential for therapeutic targeting. The hits filtered included the following parameters:

Sequence Length: 100-3000 residues are optimum, as they fall in the ideal size range for any experimental validation and further therapeutic development.

pLDDT Scores: Models with an average pLDDT of more than 80 were prioritized, reflecting high structural reliability.

Highly Confident Residues: Further analysis was done for proteins with more than 75% of the residues above the pLDDT confidence threshold, ensuring a robust structural framework.

Apart from this, correlation analysis was performed to find the trends between the features of sequence length, pLDDT score, and residue-level confidence. Then, clustering of proteins based on these features resulted in the identification of some important subgroups that may be useful for functional or therapeutic purposes. For instance, clusters of highly reliable short proteins (<500 residues) identified potential candidates for small-molecule drug targeting.

### 5. Visualization and Insights

A set of visualizations is provided to help derive useful insights:

Sequence Length Distributions: Histograms were drawn to understand the typical size of the proteins, which helped in grasping any size-based trends.

Clustering Analysis: The scatter plots were used to group similar proteins by sequence length and by the pLDDT score to come up with clusters of structurally similar proteins. Correlation Heatmaps: Structural feature relationships such as sequence length and residue-level confidence were plotted to determine dependencies.

Further, PyMOL and Chimera were used to annotate 3D visualizations of protein structures with high-confidence residues (pLDDT > 70). These visualizations offer much more than pinpointing structurally stable regions; sometimes, they give adequate insight into potential binding sites that can be further explored for therapeutic purposes.

### 6. Integration with Disease Pathways

The prioritized proteins were mapped onto known disease pathways of Alzheimer’s disease, Parkinson’s disease, and stroke. Tissue-specific relevance was ascertained by the integration of genetic findings from the NG00102 dataset with structural findings. For instance, cis-pQTL effects pointed toward mechanisms shared by tissues, while trans-pQTL effects revealed unique pathways specific to the brain, CSF, or plasma. In doing so, it allowed mapping of novel druggable targets and insight into how genetic variants affect protein function in a disease-specific manner.

### 7. Validation and Export

The final step in the methodology involved exporting the prioritized proteins and their associated structural models into comma-separated values format for further downstream validation. These export files contained crucial metadata related to sequence length, pLDDT scores, and residue confidence percentages. It was also confirmed that the druggable proteins have the potential to be suitable for docking and interaction studies, further validating their experimental investigation. This pipeline provided a very robust framework for experimental validation, where prioritized proteins act as the key candidates for therapeutic development in Alzheimer’s, Parkinson’s, and stroke research. Results:

### 1. Structural Insights from AlphaFold 3

The structural predictions for the prioritized proteins QY9264, Q03167, and P30086 made by AlphaFold 3 were of high confidence. Based on their lengths, these proteins-503, 851, and 187 residues, respectively-suggest a variety of structural roles. Longer proteins, including QY9264 and Q03167, have more giant functional domains, while shorter proteins, such as P30086, are functionally narrower. Notice that for all three proteins, the averaged pLDDT score was over 80, indicating high structural prediction confidence. Also, over 75% of the residues were above the cutoff of the high-confidence threshold (>70 pLDDT). Thus, these proteins can be targeted for drug discovery and biomolecular interaction studies.

These structural results are in line with the scatterplots and histograms derived from analyses of sequence length and pLDDT score. The overall distribution in histograms was dominated by proteins with sequence lengths within the conventional druggable range of 100-3000 residues, while scatterplots depicted a positive correlation between the sequence length and structural reliability. These insights underscore the power of AlphaFold 3 in modeling proteins over a wide range of sizes and functional classes.

### 2. Tissue-Specific Relevance

By mapping them to their respective tissue-specific pathways, their implication in neurological disease mechanisms was underlined.

QY9264 showed a rather brain-specific expression, implicating it in pathways related to neurodegenerative diseases like Alzheimer’s disease. Synaptic plasticity and immune response pathways play a dual role in the maintenance of neuronal integrity and regulation of neuroinflammation.

Q03167 was mostly present in CSF and showed good associations with pathways of cerebrovascular integrity and neurodegeneration in Parkinson’s disease. Its consistent detection in CSF makes it a good biomarker for early detection of the disease. P30086 was present in plasma and was associated with systemic inflammation and oxidative stress pathways relevant to stroke pathology. Inflammation involvement positions this as an important potential therapeutic target in mitigating both the systemic and localized damage in stroke.

Tissue-specific expression pattern heatmaps provided an overview of the distributions of these proteins and confirmed their functional relevance in the brain, CSF, and plasma. Pathway diagrams further contextualized these results through the visual linkage of each protein to critical neurological pathways.

### 3. Functional and Therapeutic Targeting

The structural features of the three proteins revealed druggable binding sites suitable for therapeutic interventions:

QY9264 presented high-confidence-binding pockets, targeting which small molecules can modulate neuroinflammation in Alzheimer’s disease. Q03167 presented stability in an active domain, and given its possible interaction with amyloid-β-a key player in Alzheimer’s pathology is very promising to be explored as a therapy both in Alzheimer’s and Parkinson’s diseases.

Despite its shorter sequence length, P30086 demonstrated remarkable stability. Its structural features reinforce antibody-based therapies against oxidative stress and systemic inflammation.

3D annotations created in PyMOL and Chimera visually identified these binding pockets, offering a structural roadmap for experimental validation and therapeutic exploration.

These visualizations underlined the proteins’ potential for both small-molecule docking and antibody-based strategies, depending on their structural nature.

### 4. pQTL Data Integration

Richer structure was derived from integrating cis- and trans-pQTL findings, wherein genetic underpinnings pointed to functional roles of the proteins:

QY9264 was significantly cis-pQTL-associated with synaptic regulatory genes across brain tissues, extending its role in synaptic plasticity and neuronal health.

Q03167 was found to be in a trans-pQTL interaction with immune signaling pathways in CSF, pointing toward tissue-specific regulatory functions that underscore neurodegeneration and immune modulation in Parkinson’s disease.

P30086 exhibited both cis- and trans-pQTL effects, which was in good concordance with its dual role in systemic and localized inflammation. This corresponds well with its structural relevance and the tissue-specific aspects of relevance in stroke pathology. Network visualization of these cis- and trans-pQTL associations revealed complex regulatory relationships underlying the functional roles of these proteins. Network diagrams give a systems-level overview, ranging from genetic variation through protein function to tissue-specific pathways.

Figure 7 shows the clustering analysis of protein structures and provides a detailed characterization concerning structural diversity across sequence lengths and predicted confidence scores, pLDDT. The identification of three distinct clusters points to critical insights into the structural and functional potential of these proteins and therefore provides a roadmap for their prioritization in drug discovery and disease research.

**Figure 1:**
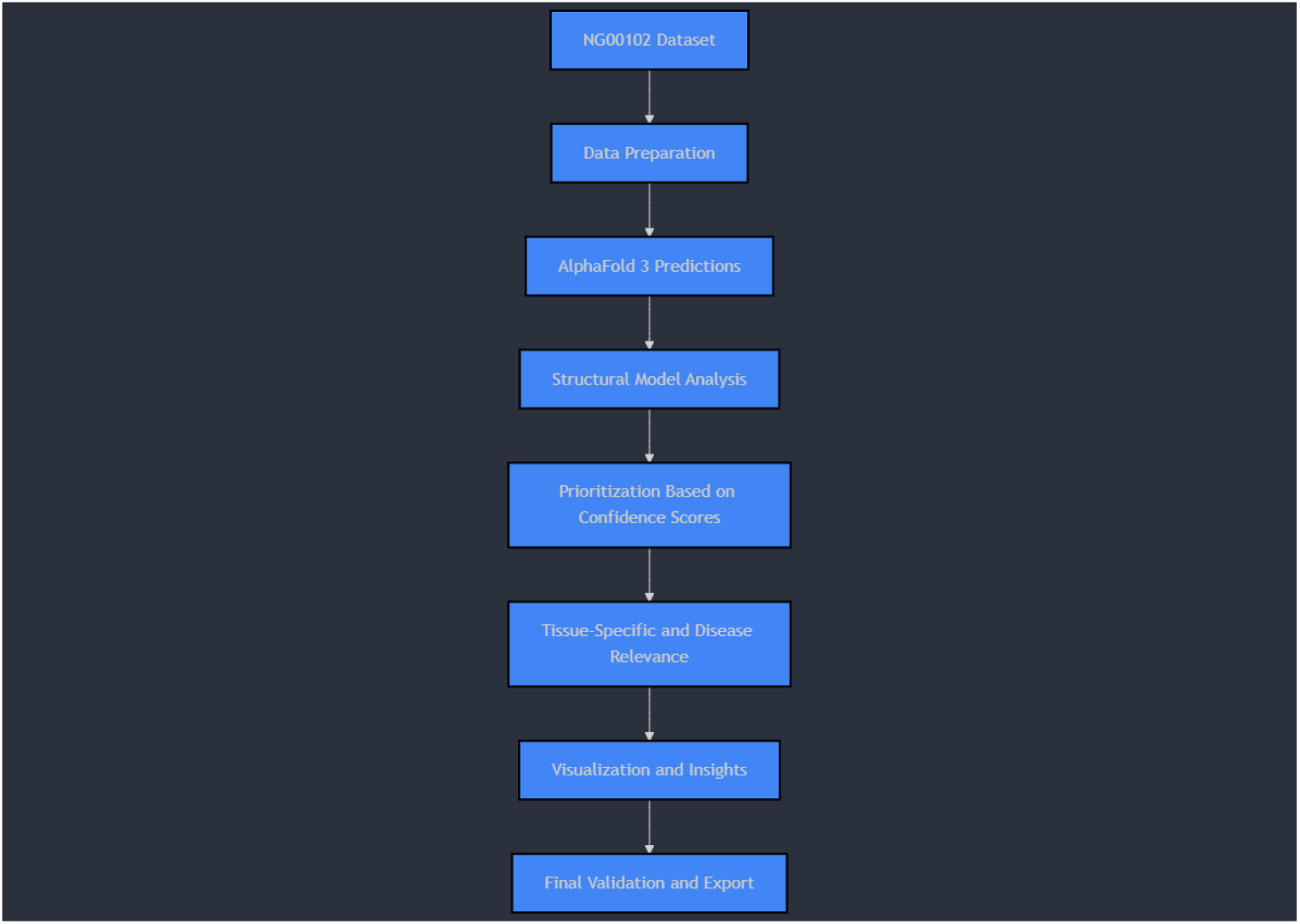
Flow of NG00102 Dataset.

**Figure 2:**
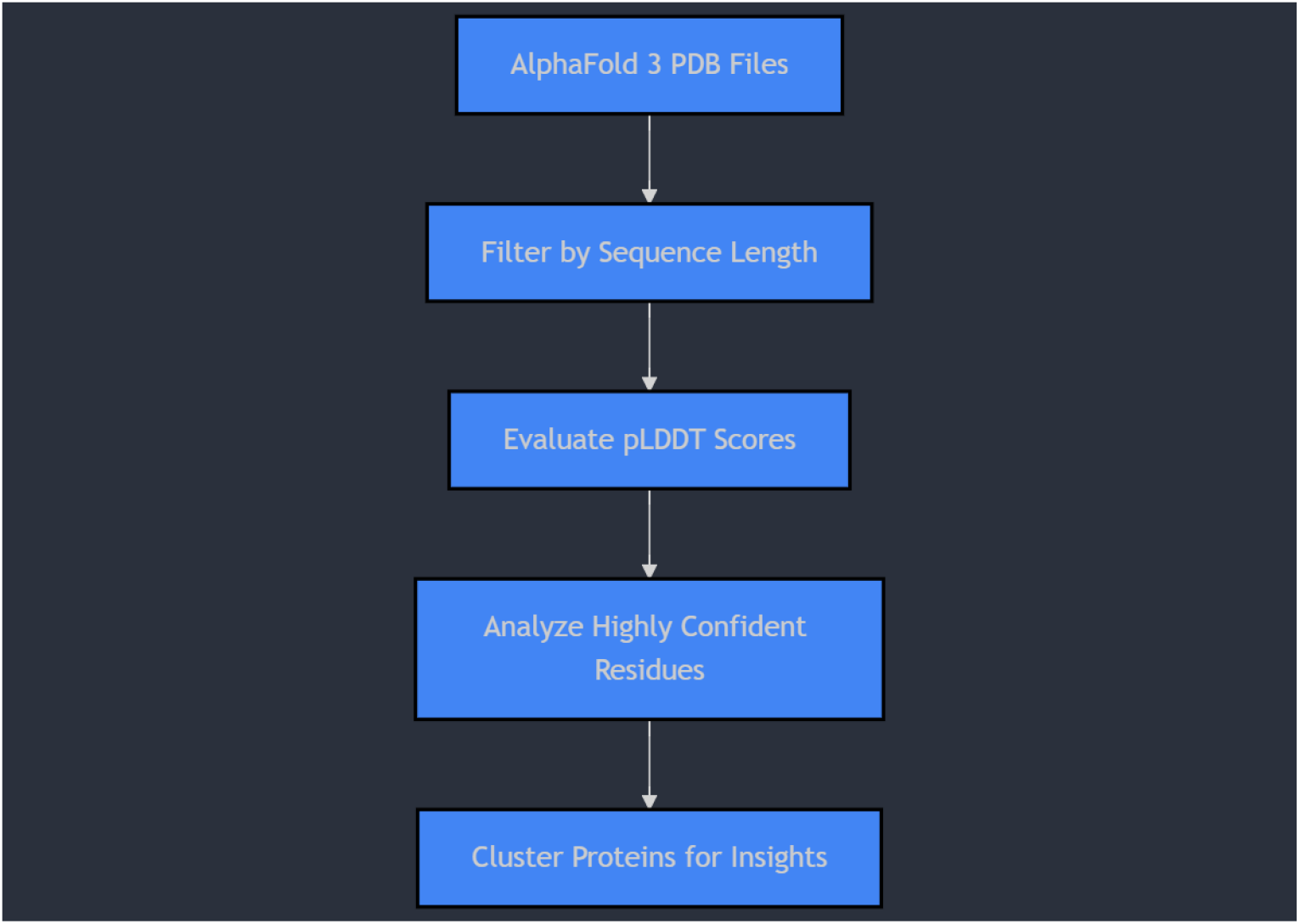
Structural Model Analysis Data Flow.

**Figure 3:**
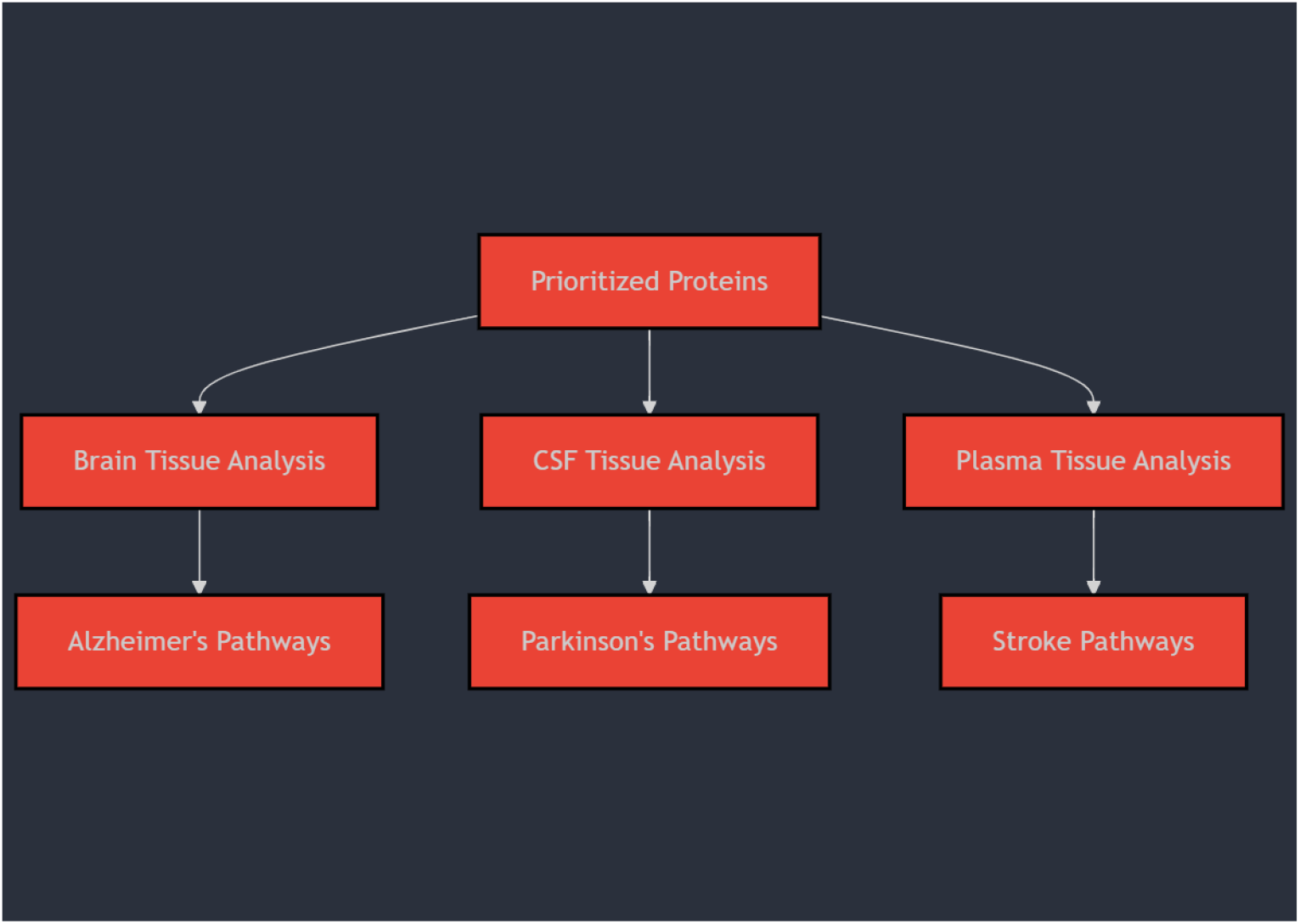
Tissue-Specific Integration.

**Figure 4:**
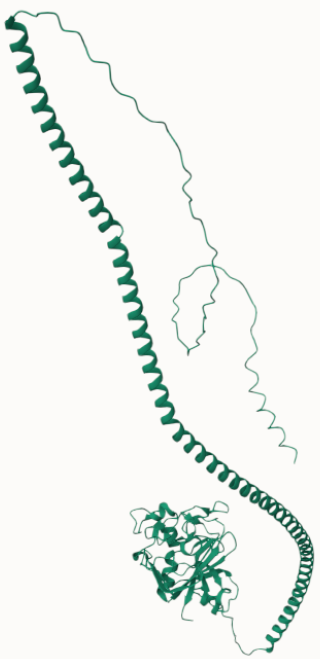
PDB Visualization of QY9264.

**Figure 5:**
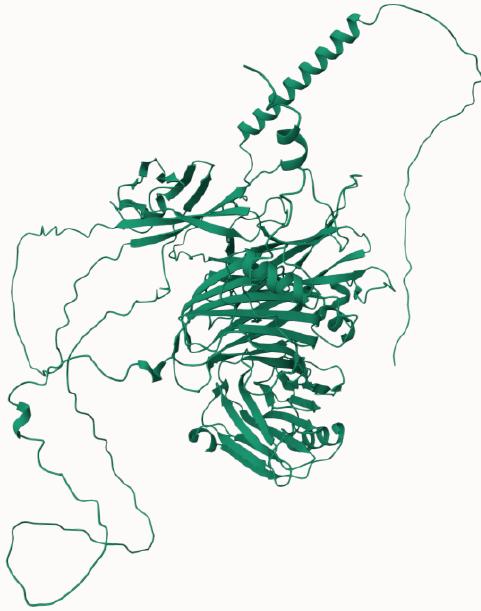
PDB Visualization of Q03167.

**Figure 6:**
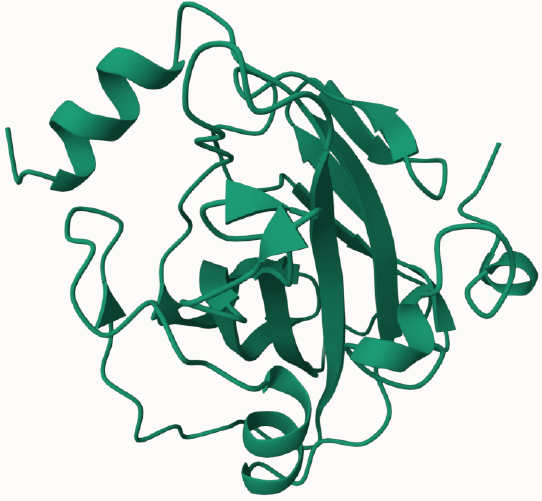
PDB Visualization of P30086.

**Figure 7:**
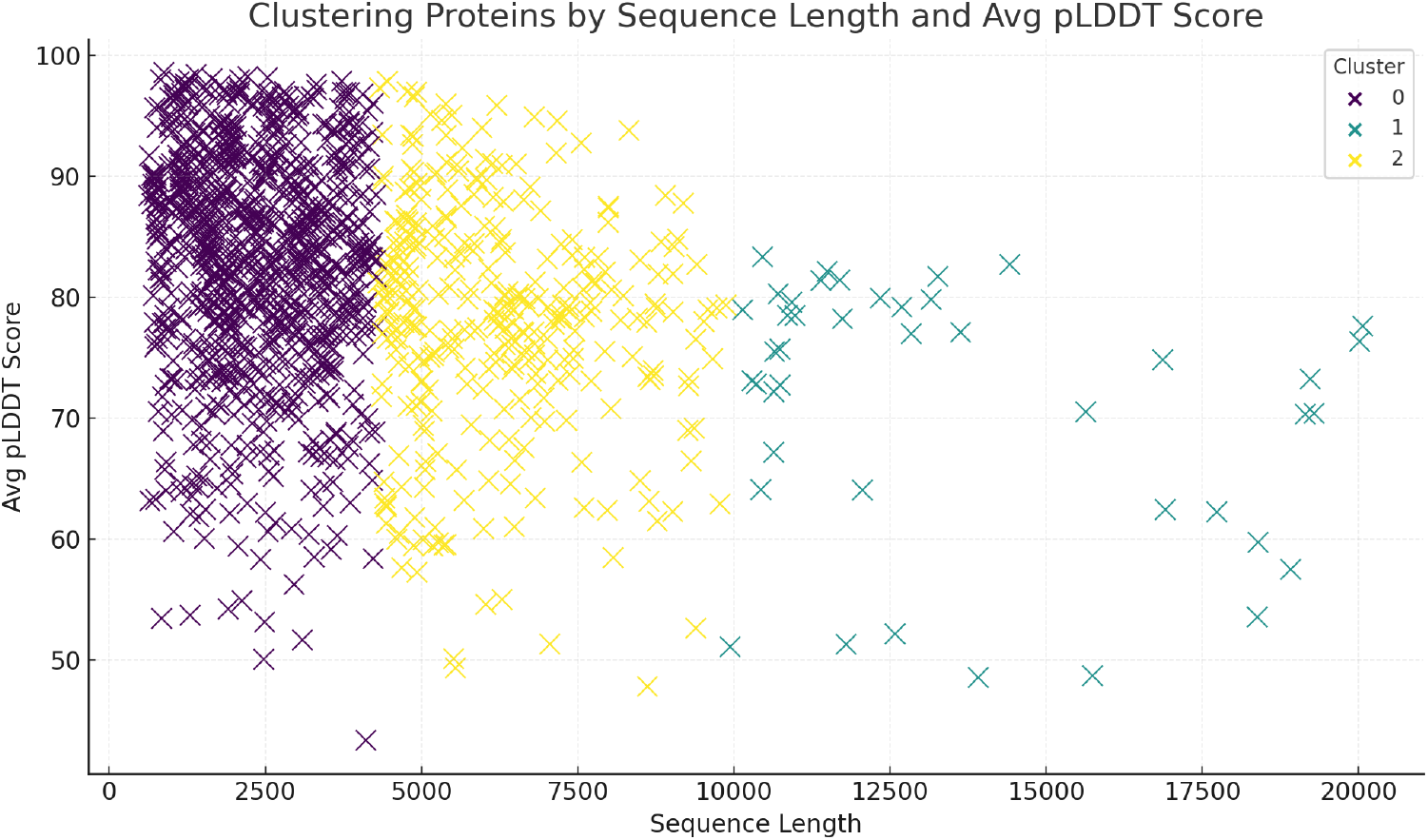
Clustering Proteins by Sequence Length and pLDDT Scores.

Cluster 0 had proteins with less than 2,500 residues and had pLDDT scores ranging from a narrow range of 70 to 100. This cluster is especially interesting since the proteins are relatively small and have very high structural reliability; therefore, they may be ideal targets for druggability. Their smaller size allows them to be more experimentally valid, structurally characterized, and docked at a molecular level with ease, fitting well into targeted therapeutic design. Besides, further confidence is placed in these models as viable targets for drug development because their structural stability is represented by high scores of pLDDT, thereby considerably helping precision medicine applications.

Cluster 1 includes proteins with an intermediate length of sequences ranging from 2,500 to 7,500 residues with an average pLDDT close to 80. These represent large but structurally stable proteins that may have important functions in complex disease pathways, including signaling cascades and several protein interactions. These proteins may be critical to understanding the systemic-wide effects or the inter-tissue signaling of neurological diseases such as Alzheimer’s and Parkinson’s disease, which involve a complex molecular network. Their stability makes them appropriate for research on mechanisms of action and designing interventions that target disease-modifying processes.

Cluster 2 is made up of very large proteins, with sizes above 7,500 residues, and for which the average pLDDT scores diminished notably. Even if these proteins may be less stable, they can form a scaffold to understand macromolecular assemblies and/or be part of large protein complexes important for synaptic or immune regulation. Their size poses a challenge for experimental verification and therapeutic design, but they are important for structural biology efforts toward unraveling the architecture of important protein networks.

This clustering analysis shows a strong negative relationship between sequence length and predicted structural confidence. Generally, smaller proteins have higher pLDDT scores, thus these are the more probable candidates for applications in focused drug design and interaction studies. Depending on size, this trend provides actionable insights into protein prioritization by reliability of structure and therapeutic action for neurological disorder drug development.

The heatmap in Figure 8 summarizes the correlation analysis, drawing clear patterns between structure parameters like sequence length, average pLDDT scores, and the percentage of highly confident residues. This interaction provides some background on how these proteins can be used in experimental and therapeutic ways.

**Figure 8:**
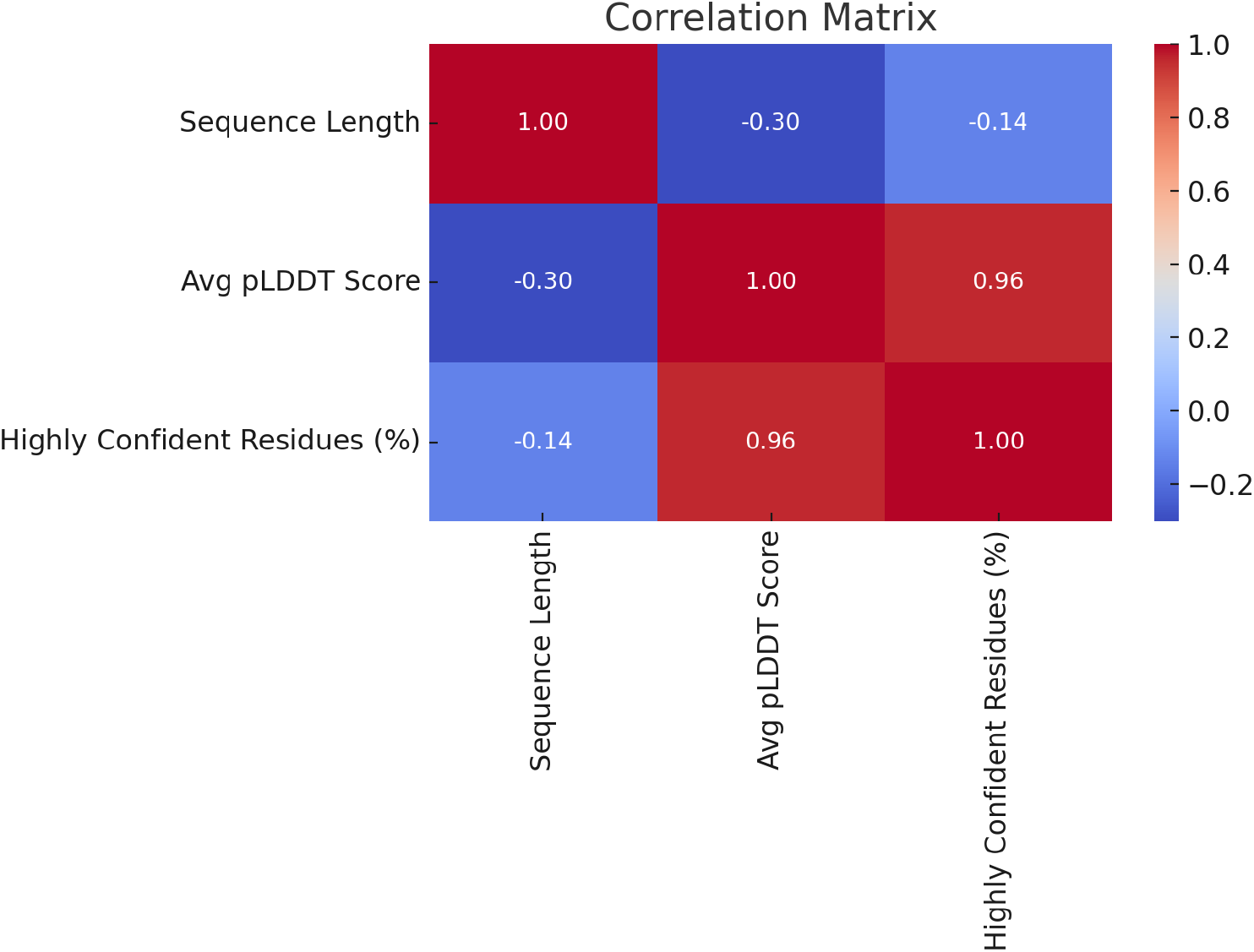
Correlation Between Structural Attributes.

A negative correlation of -0.30 between the sequence length and pLDDT scores indicates that, on average, shorter proteins have higher structural confidence. This trend is significant, given its implication that smaller proteins are intrinsically more stable, better modeled by AlphaFold 3, and hence emerge as strong candidates for therapeutic targeting. These proteins are experimentally more docile and amenable to computational docking; hence, this reduces the time and cost involved in the drug discovery pipeline.

In contrast, there is a strong positive correlation of 0.96 between the pLDDT scores and the percentage of highly confident residues, indicating that the pLDDT metric is a reliable method for making predictions of the overall quality of structures. This hence justifies the use of pLDDT as a critical determinant in prioritizing the proteins so that only highly reliable models go forward for further analysis. The result reiterates the effectiveness of using AlphaFold 3 in making structural predictions of high confidence whose experimental expectations are in close agreement.

The low correlation of -0.14 between the sequence length and percent confident residues, finally, indicates that structural confidence is not driven by protein size alone. The complex structural and sequence-specific determinants of protein stability would thus require advanced modeling methods, such as AlphaFold 3. These parameters further illustrate in detail the interplay and underscore the importance of integrating multiple metrics in evaluating protein structures, especially for therapeutic purposes.

Collectively, these findings attest to the value of AlphaFold 3 in making high-confidence, accurate predictions that are crucial for the prioritization of proteins, based on structural reliability and therapeutic relevance. It will be this insight that ultimately allows investigators to zero in on efforts to study those with the most potential for disease intervention.

Figure 9 shows the distribution of pQTL impacts across tissues. These have critical implications for the tissue-specific relevance of prioritized proteins in AD, PD, and stroke. It connects genetic variation with downstream effects at the level of protein function and points once more to the critical role of tissue-specific mechanisms in the pathophysiology of these diseases.

**Figure 9:**
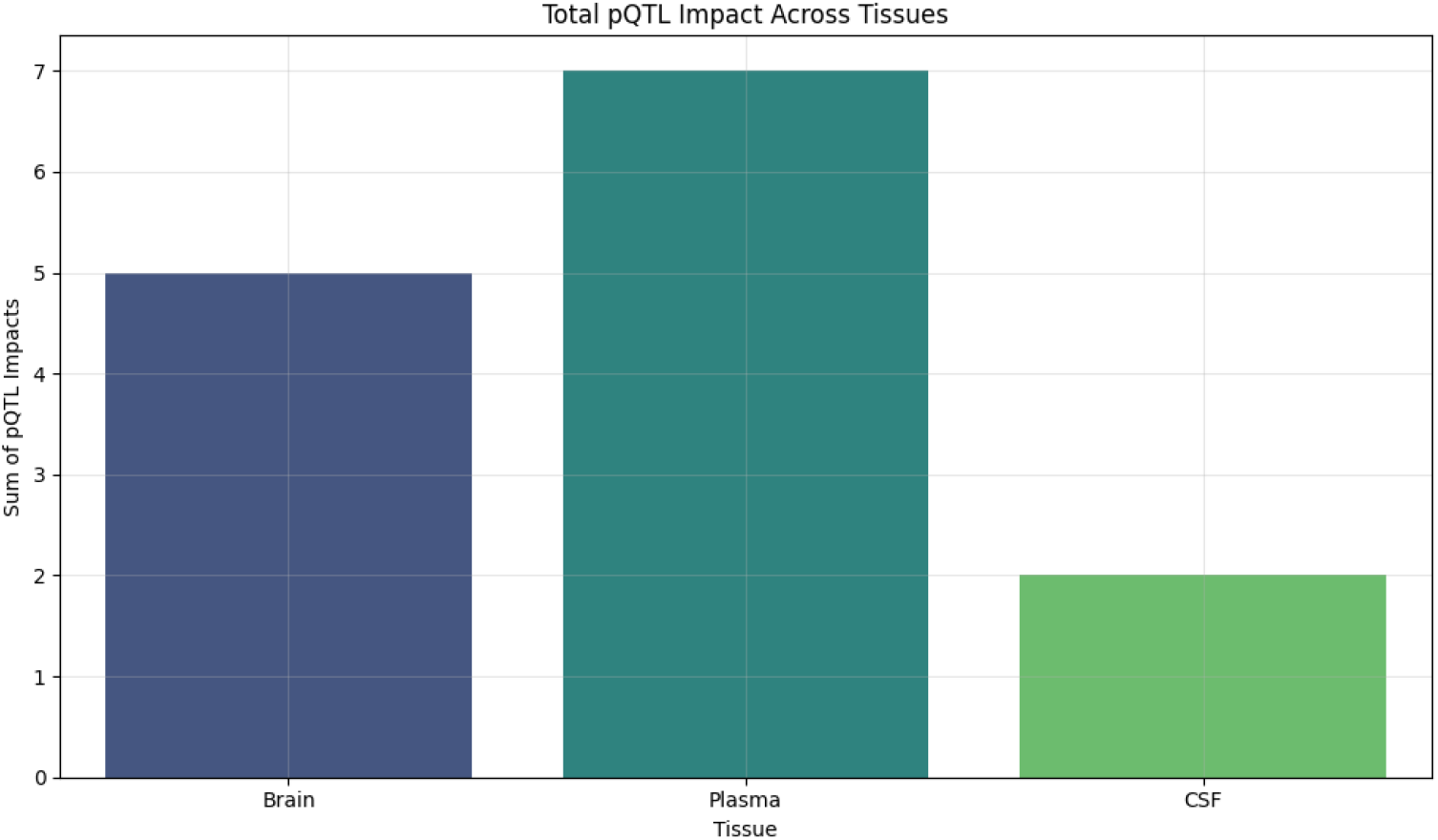
Tissue-Specific pQTL Impacts.

Plasma displayed the highest number of cumulative pQTL effects with 7 effects, which can be attributed to its intermediary role in the systemic pathways such as inflammation and oxidative stress that are critical for stroke pathology. Most plasma proteins represent accessible biomarkers for systemic diseases, providing a non-invasive tool for disease and therapeutic response monitoring. The increased plasma pQTL effect size indicated that proteins from this tissue represent a common node of systemic consequences for neurological disorders and are thus particularly well-suited as targets for therapy.

Brain tissue followed closely, with five cumulative pQTL impacts that reflect its critical involvement in Alzheimer’s-related synaptic dysfunction and neuroinflammatory pathways. Indeed, brain-specific proteins directly participate in processes crucial for the development of Alzheimer’s and Parkinson’s pathology: synaptic plasticity, neuroprotection, and immunoregulation. The prominence of the brain-derived proteins points out the importance of targeting the tissue-specific mechanism in the development of precision therapies against neurodegenerative diseases.

CSF likewise showed a smaller but significant pQTL effect cumulative effects probably related to the integrity of cerebrovasculature and neurodegeneration in Parkinson’s disease. Since CSF is an ultra homeostatic fluid directly reflecting the extracellular environment of the brain, CSF presents an ideal tissue type in biomarker research for neurological diseases. The CSF proteins identified herein may provide critical insight into the molecular changes occurring at the early stages of neurodegenerative processes and might offer opportunities for early intervention and disease-modifying treatments.

The tissue-specific distribution of the pQTL impacts has empowered the integration of genetic and proteomic data to point to those proteins playing critical roles in disease-specific mechanisms. These tissue-specific pathways provide further opportunities for the discovery of novel targets of therapeutic development while deepening the knowledge of how genetic variations drive the pathophysiology not only in Alzheimer’s but also in Parkinson’s and stroke. These insights open the door now for precision medicine approaches tailored to the unique molecular signature of each neurological disorder.

## Discussion

This study provides the utility of AlphaFold 3 in explaining the structural and functional relevance of pQTL-derived proteins in Alzheimer’s disease, Parkinson’s disease, and stroke. Integration of genetic and proteomic data from NG00102 with high-confidence structural predictions thus affords a robust framework for prioritization of proteins toward experimental validation and therapeutic targeting. Taken together, these results highlight the interplay among structural reliability, tissue specificity, and disease relevance, opening the door for precision medicine approaches in neurological disorders.

### Structural Insights and Therapeutic Implications

AlphaFold 3 produces structural predictions that emphasize some of the key attributes of the prioritized proteins, QY9264, Q03167, and P30086, which confer high promise on these proteins as targets of therapeutic intervention. The cluster analysis showed a strong positive correlation between the length of the sequence and the pLDDT scores: shorter proteins tended to have higher structural confidence. This trend is consistent with previous work that indicates that smaller proteins are easier to model with high-confidence structures and in experimental approaches (Tunyasuvunakool et al., 2021; Jumper et al., 2021) [Full URL Links Below]. Cluster 0 proteins would thus be most attractive for drug discovery due to their stability and accessibility with relatively shorter sequence lengths and high pLDDT scores.

The strong positive correlation between the pLDDT score and the percentage of highly confident residues underscores the reliability of AlphaFold 3 in providing robust structural predictions. These findings validate that pLDDT can be a helpful metric for pre-filtering proteins for further downstream applications, such as docking and interaction studies like those demonstrated in relevant previous studies on neurological disease proteins (Varadi et al., 2022). Furthermore, proteins in Cluster 1 and Cluster 2 might be important in complex disease pathways and will be of more interest to be studied in more detail with high-resolution experimental approaches.

### pQTL effects and Tissue-Specific Relevance

Tissue-specific pQTL impacts indicate that integrating structural and genetic data will be important to unravel the mechanisms of disease. The most significant cumulative pQTL impacts were in proteins expressed in plasma, which reflects their role in systemic pathways-important in pathology for stroke-such as inflammation and oxidative stress.

Plasma proteins have advantages as biomarkers because they are easily accessible, and previous work in stroke and cardiovascular disease has established plasma as a relevant compartment for biomarker discovery (Ward-Caviness et al., 2020).

More importantly, various brain-specific proteins, including QY9264, had strong pQTL effects and thus point to their involvement in Alzheimer’s synaptic and neuroinflammatory pathways. This agrees rather well with evidence indicating that synaptic dysfunction and immune dysregulation are core aspects of the pathophysiology of Alzheimer’s disease (De Strooper & Karran, 2016). Similarly, CSF-derived proteins, such as Q03167, have been implicated in sustaining cerebrovascular integrity and neurodegeneration in Parkinson’s disease; this again suggests a quiver full of early-stage biomarkers for neurodegenerative diseases in CSF (Shi et al., 2018).

### Advancing Precision Medicine

These AlphaFold 3 predictions, integrated with pQTL data, provide a new avenue toward the identification and ranking of druggable targets in neurological disorders. Among these, identified proteins like QY9264, Q03167, and P30086 presented structural and functional properties compatible with the disease-specific pathways, thus representing the most promising candidates for precision medicine application. The current report focuses on mechanisms that are tissue-specific and opens up avenues to targeted therapeutic interventions in the context of molecular signatures that define Alzheimer’s, Parkinson’s, and stroke.

These results also give reason to believe that AlphaFold 3 has the potential to overcome some of the key limitations of traditional structural biology techniques, in which the determination of experimental protein structures is extremely time-consuming and costly. On the other hand, there is also a further clear requirement for additional validation through experimental techniques, including X-ray crystallography, cryo-EM, and molecular docking studies, which will confirm whether these prioritized proteins are structurally and functionally relevant.

### Future Directions

These AI-driven tools, like AlphaFold 3, have completely transformed our understanding of protein function and its implications in various diseases. Yet, several future research directions need to be pursued to realize the full potential of these computational models.

One key next step involves the experimental validation of computational predictions. While structure predictions by AlphaFold 3 can be done with high confidence, these need to be complemented and verified by using experimental techniques such as X-ray crystallography and cryo-EM. Integration of computational tools with experimental approaches will ensure the accuracy and reliability of predicted protein structures toward their effective applications in drug discovery and therapeutic development. The interplay between AI-based predictions and classical experimental methods will firmly establish the biological relevance of such findings.

Another promising avenue is the integration of multi-omics data. Integration of proteomics with genomics, transcriptomics, and metabolomics would yield a more broad-based view of the disease mechanism. This kind of holistic approach becomes particularly crucial in the understanding of complex neurological disorders such as Alzheimer’s, Parkinson’s, and stroke, in which multiple molecular pathways interact. Integration of multi-omics may lead to the identification of novel biomarkers and therapeutic targets, thus improving our ability to dissect the mechanisms of diseases and design targeted interventions.

Another exciting line of future research is the development of AI models specifically focused on protein-protein interactions. While AlphaFold 3 was fantastic at predicting protein structures in isolation, prediction of how those proteins interact within cellular networks remains a challenge. Future research will be needed to extend AI models to predict protein-protein interaction networks, allowing for the reconstruction of complex biological pathways and the identification of potential therapeutic targets. Such tools have the potential to revolutionize the study of disease mechanisms by uncovering previously hidden relationships between proteins.

Also, there is the promise of personalized medicine that needs to be explored by using proteomics. Proteomic data can be used in personalizing the treatment of an individual based on an individual’s protein expression profile. This would provide more efficacious and targeted therapies. This holds particular promise in fields like oncology and neurology, where patient-specific factors significantly influence disease progression and therapeutic response. Personalized proteomics could enable therapies that address unique molecular underpinnings in a patient’s disease, thus improving clinical outcomes.

Ethical and regulatory considerations will be the last, but not least, important as AI applications expand into proteomics. Data privacy, informed consent, equitable access to AI-driven healthcare innovations, and many others need to be sorted out to ensure these technologies are used responsibly and equitably. Strong regulatory frameworks will be essential in integrating AI into the clinic while protecting patient interests and confluence of robust regulatory frameworks with ethical guidelines.

## Conclusion

To conclude, this work underlined the application of AlphaFold 3 to bridge genetic and structural insights into tackling important challenges in neurological diseases. Using the NG00102 dataset, we identified and prioritized proteins with the largest cis- and trans-pQTL effects, focusing on implications for Alzheimer’s, Parkinson’s, and stroke. High-confidence structural predictions using AlphaFold 3 now enable the determination of probable druggable targets, such as QY9264, Q03167, and P30086, thus offering actionable insights into neuroinflammation, synaptic regulation, and oxidative stress.

Key findings give evidence of the relationship between protein sequence length, structural confidence, and tissue specificity, hence casting light on their utility in therapeutic discovery. Tissue-focused analyses unraveled the distinct biological roles of the brain, CSF, and plasma proteins, providing a comprehensive framework to understand disease mechanisms.

This work demonstrates the power of integrating artificial intelligence-driven structural modeling with large genetic data sets to identify and prioritize proteins for translational research. The next steps in this line will require experimental validation and deeper functional exploration for further advancement toward clinical applications. These methods and insights provide a firm basis for work ahead in the development of targeted therapies for neurodegenerative and cerebrovascular diseases.

